# Membrane Targeted Azobenzene Drives Optical Modulation of Bacterial Membrane Potential

**DOI:** 10.1101/2022.09.05.506195

**Authors:** Tailise Carlina de Souza-Guerreiro, Gaia Bondelli, Iago Grobas, Stefano Donini, Valentina Sesti, Chiara Bertarelli, Guglielmo Lanzani, Munehiro Asally, Giuseppe Maria Paternò

## Abstract

Recent studies have shown that bacterial membrane potential is dynamic and plays signalling roles. Yet, little is still known about the mechanisms of bacterial membrane potential regulation –owing in part to a scarcity of appropriate research tools. Optical modulation of bacterial membrane potential could fill this gap and provide a new approach to studying and controlling bacterial physiology and electrical signalling. Here, we show that a membrane-targeted azobenzene (*Ziapin2*) can be used to photo-modulate the membrane potential in cells of the Gram-positive bacterium *Bacillus subtilis*. We found that upon exposure to blue-green light (λ = 470 nm), isomerization of *Ziapin2* in the bacteria membrane induces hyperpolarisation of the potential. In order to investigate the origin of this phenomenon we examined ion-channel-deletion strains and ion channel blockers. We found that in presence of the chloride channel blocker idanyloxyacetic acid-94 (IAA-94) or in absence of KtrAB potassium transporter, the hyperpolarisation response is attenuated. These results reveal that the *Ziapin2* isomerization can induce ion channel opening in the bacterial membrane, and suggest that *Ziapin2* can be used for studying and controlling bacterial electrical signalling. This new optical tool can contribute to better understand microbial phenomena, such as biofilm electric signalling and antimicrobial resistance.

## Introduction

Genetic and non-genetic optomodulation is recognised as a transformative technology in neuroscience^[1–4]^. For example, optogenetics has been successful in bidirectionally controlling animal behaviours^[5,6]^, and it has a foundation for treating neuropsychiatric disorders and rebuilding vision^[7–9]^. Non-genetic optomodulation is expected to broaden the scope of applications and complement genetic approaches as it can mitigate some deep concerns associated with real-life applications of genetically modified organisms.

Recently, we have introduced a molecular optomechanical light transducer, named *Ziapin2*, which is able to drive optical modulation of the electrical properties of membranes in primary culture neurons and *in vivo* mouse brain^[10]^. Specifically, *Ziapin2* is an amphiphilic azobenzene with a strong non-covalent affinity to the plasma membrane^[10,11]^ (Figure 1). Its optomodulation ability resides in the fact that the dark-adapted *trans* isomer causes a thinning of the lipid bilayer via a dimerization mechanism, while illumination with visible light (~470 nm) leads to a membrane relaxation that follows disruption of the azobenzene dimers (Figure 1). Consequently, this brings about a light-driven decrease of the membrane capacitance and causes transient hyperpolarisation. Importantly, it was demonstrated that *Ziapin2* is nontoxic to neurons and can be used to activate cortical networks when injected into the mouse somatosensory cortex^[10]^.

**Figure 1.**
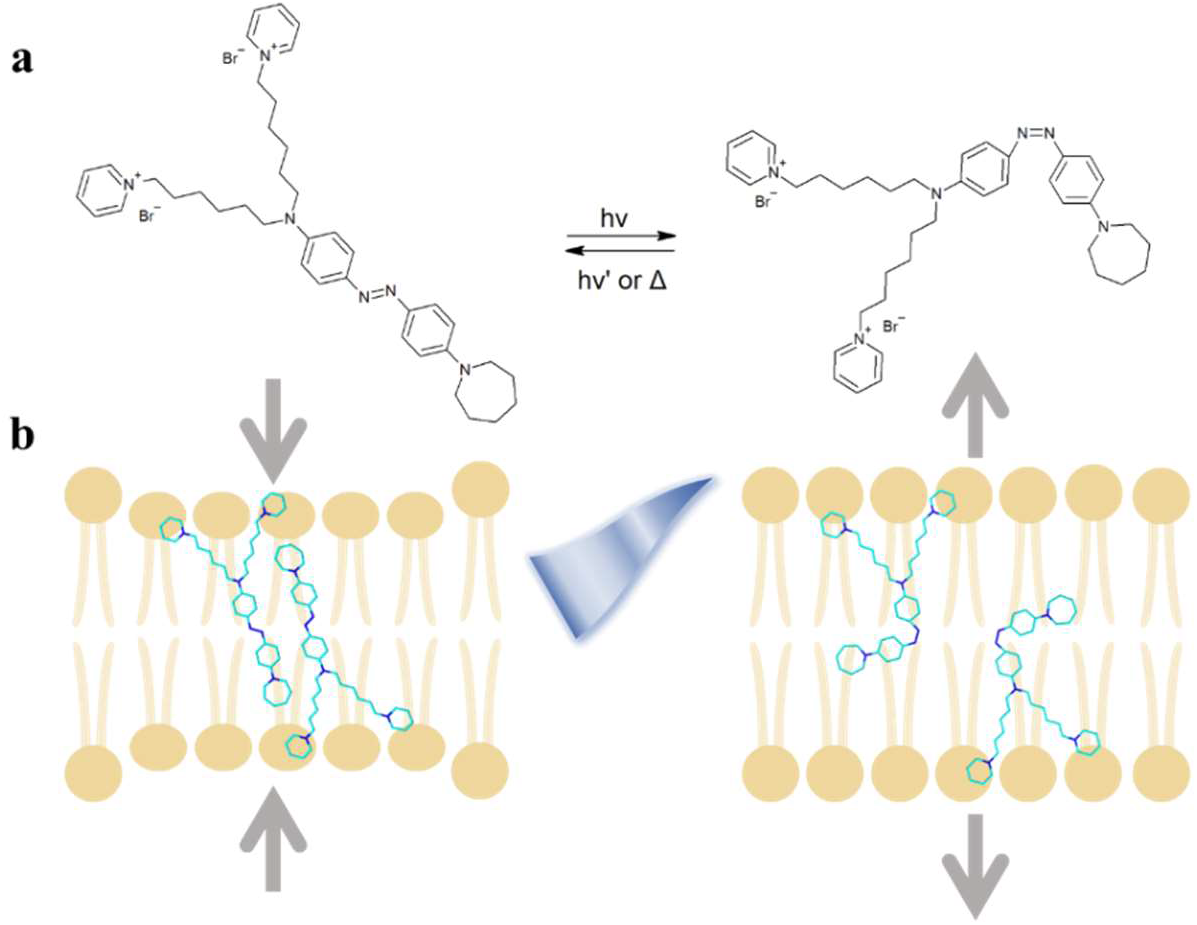
Illustrative diagram of photo-induced *Ziapin2* isomerisation. a) Molecular structure of *Ziapin2* and representation of its isomerization reaction. b) The optomechanical action of *Ziapin2* when sitting in the lipid membrane. In the *trans* elongated form, *Ziapin2* is able to dimerise within the lipid membrane, leading to a decrease in the thickness and an increase in the membrane capacitance. On the other side, illumination with cyan light (470 nm) triggers *Ziapin2* isomerisation into its cis bent form, an effect that disrupts the dimers and leads to an increase in the thickness and a decrease of the membrane capacitance.^[10,11,25–27]^

The mechanism of action of *Ziapin2* optomodulation suggests that, in principle, it may be possible to use for controlling the membrane potential of non-animal cells –such as bacteria. This possibility is intriguing in the light of recent discoveries that bacterial membrane potential can exhibit neuron-like spiking and oscillatory dynamics^[12–14]^. More specifically, spiking membrane potential dynamics in *E. coli* has been shown to play a role in mechanosensation^[15]^. The oscillatory dynamics of *B. subtilis* coordinate glutamate metabolism^[13]^ and allows nutrient time-sharing between colonies^[16]^, multi-species biofilm formation^[17]^ and collective antibiotic tolerance^[18]^. The membrane potential is also tied to spore formation^[19]^ and cellular responses to ribosome-targeting antibiotics^[20,21]^. These findings argue that modulating the bacterial membrane potential could provide a novel approach for controlling various membrane-potential-associated cellular processes – such as biofilm formation and antibiotic tolerance/resistance. Within this context, we recently showed that bacterial membrane potential can be altered by an externally applied electric field^[22,23]^. Optostimulation holds the potential to overcome the limitations of the electrode-based techniques, which are in general poorly suited for bacteria due to their high cell-to-cell heterogeneity, small sizes, thick cell wall and motility. In particular, optical technologies can permit to elicit and monitor signalling rapidly, remotely, and with high spatiotemporal precision. Therefore, optomodulation may be a useful tool for both basic and applied research into bacterial cell electrophysiology and bacterial electrical signalling.^[24]^

In this paper, we investigate the possibility to extend the use of *Ziapin2* to bacteria, as the translation from neurons to bacteria is not at all obvious given the very different nature of the bacteria membrane and physiology. By fluorescence time-lapse microscopy, we demonstrate the optical modulation of bacterial membrane potential driven by visible light illumination using the Gram-positive bacterium *B. subtilis* as model organism. We show that *Ziapin2* associates with B. *subtilis* membrane and can trigger a hyperpolarisation following optical stimulation. Intriguingly, the optomodulation experiments enable to unveil the involvement of KtrAB potassium transporter and uncharacterised chloride channel in the hyperpolarisation response. Our findings not only provide the proof of concept for the optical modulation of bacterial membrane potential using a photoswitching molecule but also suggest the existence of an electrical signalling cascade that can be triggered by a transient change in membrane capacitance.

## Results

### *Ziapin2* associates with the plasma membrane in *B. subtilis*

To explore whether *Ziapin2* can be used to modulate bacterial membrane potential with light, we began by examining the association of *Ziapin2* with cells. *B. subtilis* cells were incubated with 5 and 10 μg/mL *Ziapin2* in dark and under 470-nm light. First, we measured the ζ potential of cells by their electrophoretic mobility^[28,29^]. The ζ potential is the electrical potential at a colloid particle slipping plane, consisting in the interface separating mobile fluid from the fluid that remains attached to the particle surface. It is thus expected that when the positively charged *Ziapin2* is associated with the bacterial membrane, the overall negative surface potential of the cell should become less negative. Our measurements indeed show a linear rise in ζ potential with increasing *Ziapin2* concentrations, indicating the association of *Ziapin2* with the surface of *B. subtilis* cells (Figure 2a, b).

**Figure 2.**
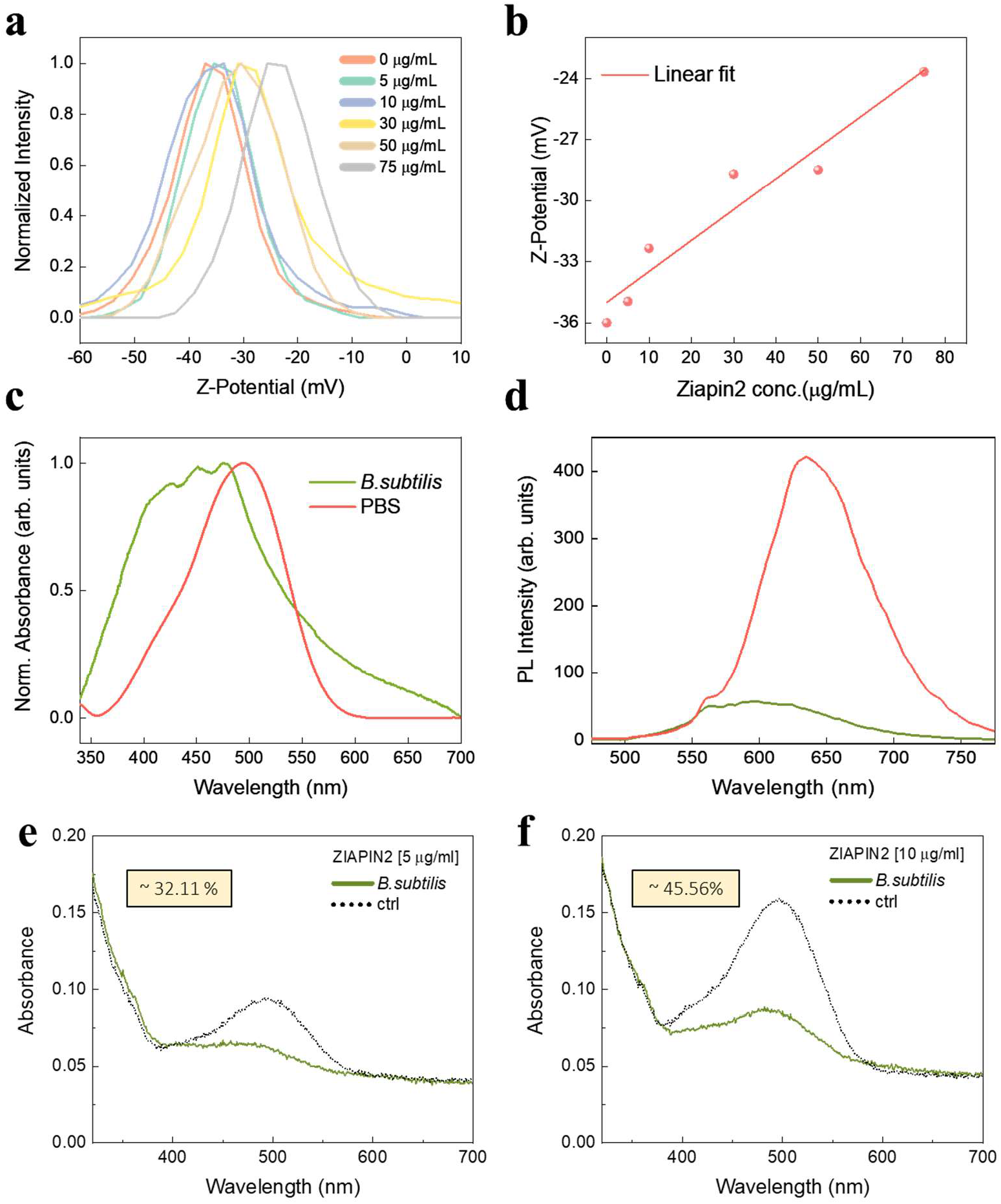
*Ziapin2* can associate with *B. subtilis* membrane. a) Variation of the distribution of ζ potential of *B. subtilis* cells as a function of *Ziapin2* concentration. b) Linear trend of ζ potential as a function of *Ziapin2* concentration. c) UV-Vis and d) PL spectra of 10 μg/mL *Ziapin2* in PBS (red lines) and in *B. subtilis* cells (green lines). PL spectra were normalized to both lamp intensity and ground state absorption, to obtain a relative PL quantum yield among the two samples. Cellular uptake experiments performed for 0.5 and 10 μg/mL of *Ziapin2*, in the supernatant (dashed line) and in the cell fraction (continuous line). See Figure S1 for the comparison between dark and light conditions.

Partitioning of *Ziapin2* into the bacterial membrane was further supported by UV-Vis and photoluminescence spectroscopies, as it happens for eukaryotic cells^[10,26]^. Specifically, the absorption spectrum of *Ziapin2* in bacteria displays a better resolved vibronic progression and a broader linewidth in comparison to *Ziapin2* in phosphate buffer saline (PBS) (Figure 2c), an effect that has been attributed to H-aggregation of the chromophore inside the lipid membrane and can be linked to *Ziapin2* dimerization at this location.^[11,30,31]^ Photoluminescence (PL) is more sensitive to the local environment than absorption as emission occurs after re-equilibration within the solvent cage and, indeed, shows clear changes in both spectral position and relative emission quantum yield. In particular, in PBS we observe both an almost 8-fold increase of the relative quantum yield and a marked red-shift (40 nm) in comparison to *Ziapin2* PL in bacteria (Figure 2d). The enhanced and red-shifted PL can be linked to the suppression of the isomerisation ability in water owing to the formation of excimer aggregates, while the membrane environment protects *Ziapin2* isomerisation. Since this is an efficient non-radiative deactivation pathway^[11]^, *Ziapin2* exhibits a relatively low emission when sitting in the membrane. Finally, the measurements of UV-vis absorption for cell fraction and supernatant showed that *B. subtilis* cells retain ~25% and ~45% of *Ziapin2* at 5 and 10 μg/mL, respectively (Figures 2e, f). No significant difference was observed between dark and 470-nm light conditions (Figure S1). These results suggest that *Ziapin2* association is not affected by the isomerization reaction and, hence, the photoreaction may be used for altering the membrane capacitance by light.

### *Ziapin2* can undergo photo-isomerisation in the bacterial membrane

To test whether *Ziapin2* can undergo light-induced isomerisation while embedded in the bacterial membrane, we employed both steady state and time-resolved photoluminescence spectroscopy. In particular, we acquired excitation/emission maps to reconstruct the *Ziapin2* deactivation scenario upon photoexcitation. The Vavilov-Kasha rule is fulfilled when the excitation profile and the absorption spectrum overlap; after absorption, the molecule relaxes to the lower excited state before emission occurs. If the two curves have different shapes, it indicates that the branching ratio between radiative and non-radiative decay paths varies with wavelength. As a test bench, we collected the PL excitation profile in DMSO, which is the solvent of choice for *Ziapin2*. Here, we observed the signature of emission from the *cis* isomer, namely an excitation peak at 370 nm, (Figure 3a)^[11]^. The *cis* isomer peak, on the other hand, was barely visible in PBS (Figure 3b), with the *trans* conformer peak at 500 nm taking precedence. This result implies that the isomerisation of *Ziapin2* in PBS is hampered, resulting in radiative deactivation within the *trans* manifold. Intriguingly, both the *cis* and *trans* isomer peaks coexisted in *B. subtilis* suspension (Figure 3c). This suggests that the bacterial membrane’s physicochemical environment restores at least partially the isomerisation ability of *Ziapin2*. We also carried out time-resolved PL experiments (Figure 3d). While the decay in PBS was mono-exponential (*τ*_1_ = 40 *ps*), the decay *in B*. subtilis cells was bi-exponential with the first component lifetime (*τ*_1_~12 *ps*), consistent with *Ziapin2* isomerisation in artificial and natural membranes^[10,11]^. All together, these data provide strong evidence for *Ziapin2* isomerisation in the bacterial membrane.

**Figure 3.**
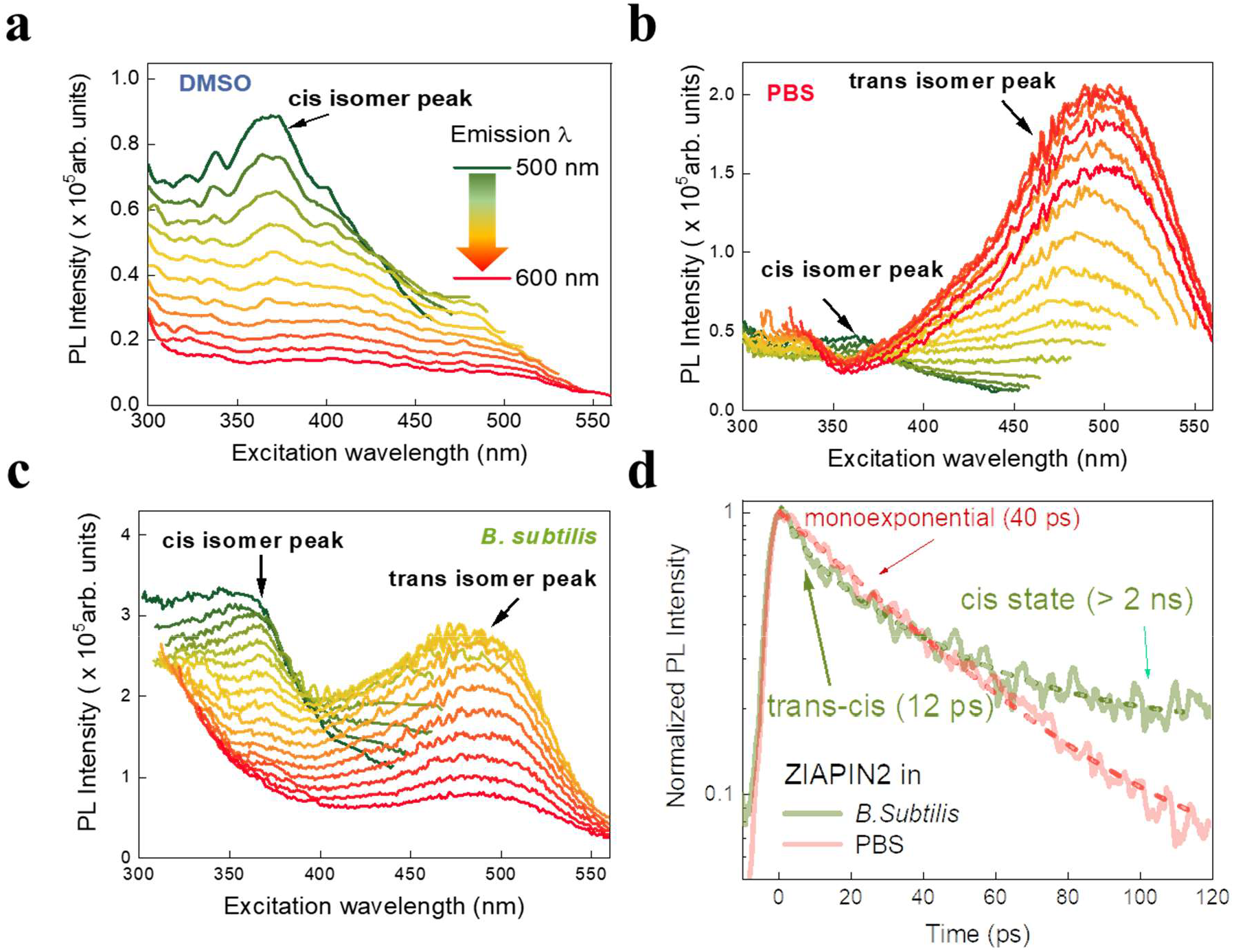
*Ziapin2* can undergo isomerisation while in bacterial membrane. Excitation–emission profiles of *Ziapin2* (10 μg/mL) in a) DMSO, b) PBS and c) *B. subtilis* cells. For each curve in plots a–c the emission wavelength is fixed at a value between 500 and 600 nm, with 10 nm steps. d) Time-resolved PL decay curves of *Ziapin2* in PBS (red line) and *B. subtilis* cells (green line). The dashed lines represent the exponential best-fit for the two curves.

### Light induces a transient hyperpolarisation in *Ziapin2*-treated bacteria

Given these results, we examined the capability of *Ziapin2* to evoke membrane potential dynamics in bacterial cells^[10]^. This would be the first translation of our non-genetic optical stimulation approach into the prokaryotic realm. First, we evaluated the cell viability upon administration of *Ziapin2* via plate reader assay, which showed that *Ziapin2* has no significant effect on cell growth when used at < 2.5 μg/mL (Figure S2). Then we proceed to study bacterial membrane potential by epifluorescence time-lapse microscopy using an optical probe, Tetramethyl rhodamine methyl ester (TMRM). TMRM is a lipophilic cationic dye that accumulates in cells with more negative membrane^[32]^. The fluorescence measurements were used to calculate the membrane potential change (ΔVm) from the resting potential (see methods). In the absence of 470-nm light stimulation (negative control), TMRM signal was stable over the course of our time-lapse experiment, regardless of the presence or the absence of *Ziapin2* (Figure 4a). We then performed time-lapse microscopy where cells were stimulated by 470 nm light for 10 sec in presence of *Ziapin2*. We confirmed that a 470-nm light stimulation does not cause a significant change in TMRM signal when *Ziapin2* is not present (Figure 4b, left). In the presence of *Ziapin2*, we observed a rise in TMRM signal following light stimulation, suggesting a hyperpolarisation by ~15 mV (Figures 4b and S3, also see Movie 1). Figure 4c illustrates the TMRM dynamics of a representative cell before and after light stimulation. TMRM signal is stable before photo stimulation, which then undergo a photo-induced hyperpolarisation followed by a gradual rebound (Figure 4c). Varying the intensities of 470-nm light, we found that the light intensity >2 mW/mm^2^ could be sufficient to cause a hyperpolarisation response (Figure S4). These results demonstrate, for the first time, that a photo-switch *Ziapin2* can indeed be used to modulate the bacterial membrane potential using light.

**Figure 4.**
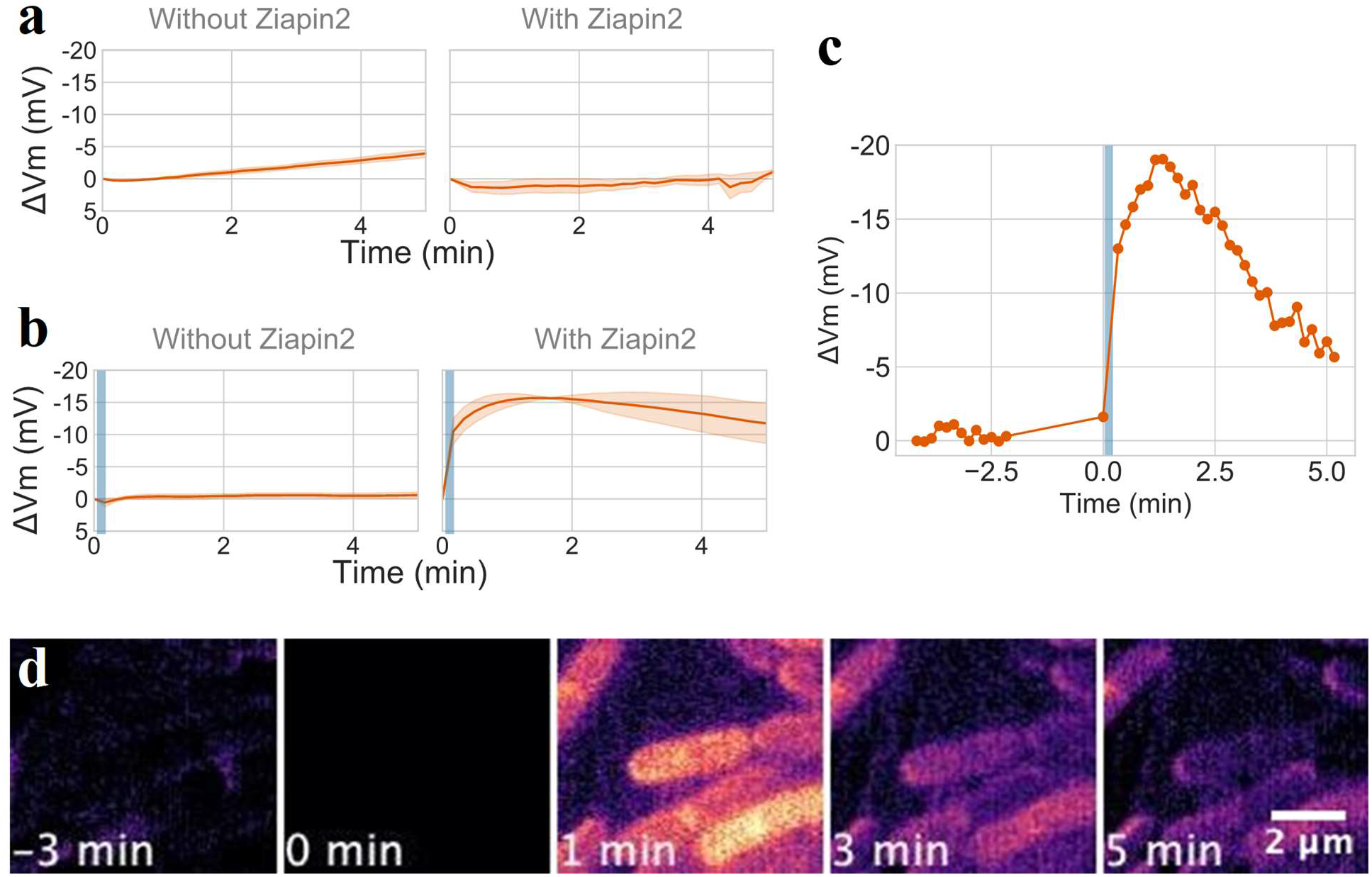
*Ziapin2* modulation of *B. subtilis* membrane potential depends on 470 nm light stimulation. a-c) Membrane potential change (ΔVm) over time, measured by TMRM fluorescence. See methods regarding the conversion of TMRM fluorescence into millivolt. The origin of time was chosen as immediately before light stimulation. The fluorescence at time 0 was used as the resting potential. Mean trace; a) without light stimulation without (left) and with (right) *Ziapin2;* b) with 10 second light stimulation (light blue) without (left) and with (right) *Ziapin2*. Shaded Areas are standard error of mean from 3 biological repeats. Blue horizontal line indicates the timing and duration of 470-nm light stimulation (20 mW/mm^2^). c) Representative single-cell time-trace of Ziapin-induced membrane potential dynamics before and after 470 nm light stimulation. d) Film strip images of TMRM signal with cells with *Ziapin2*. Cells were stimulated for 10 sec by light immediately after at time 0.

### Light-induced *Ziapin2* isomerisation leads to the opening of potassium and chloride channels

The photo-induced hyperpolarisation in bacterial cells lasted for several minutes (Figure 4). This finding is puzzling because *Ziapin2* single isomerisation event occurs in the picosecond time regime and reaches a cis-enriched photostationary state within ~ 20 seconds, while the *cis→trans* relaxation usually happens in less than one minutet^[10,11,25]^. This orders-of-magnitude discrepancy could be accounted for by a slower bioelectrical response that is triggered by *Ziapin2* isomerisation. More specifically, we hypothesised that *Ziapin2* isomerisation trigger opening of ion channels on bacterial membrane, which result in a transient hyperpolarisation.

If the light-induced hyperpolarisation is a result of biological ion channel dynamics, one would expect the response dynamics depends on the culture conditions, in particular the ones that impact the opening of ion channels. To this end, we focused on glutamate because it is known to play a central role in biofilm electrical signalling by gating the YugO potassium channel ^[13,18]^. Cells were cultured in the media with and without glutamate and examined by time-lapse fluorescence microscopy. This experiment showed that light stimulation causes a weaker hyperpolarisation response with cells in the media without glutamate (Figure 5a). This data supports the hypothesis that the photoinduced membrane potential dynamics involves a biological process.

**Figure 5.**
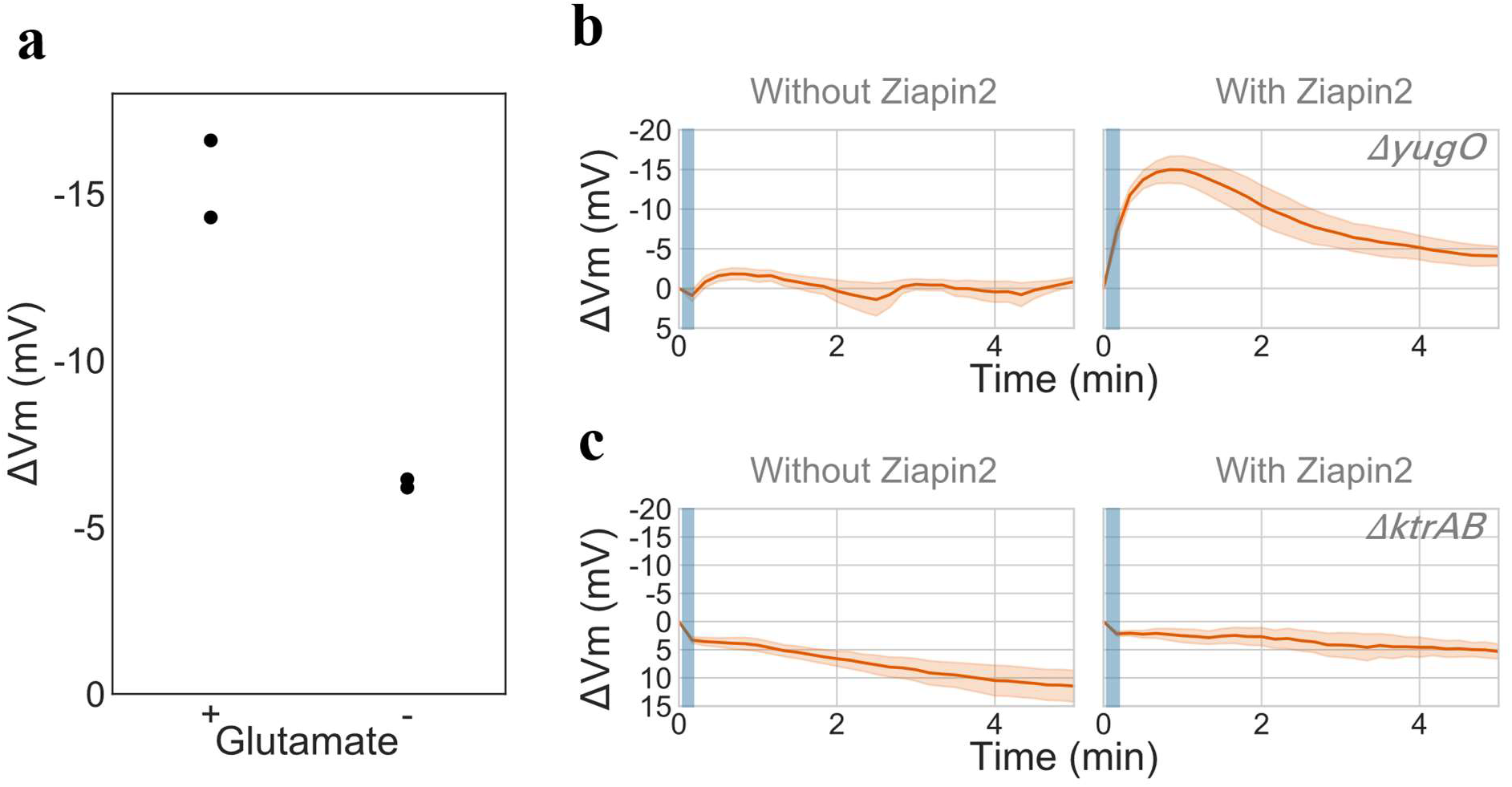
Photo-induced hyperpolarisation response depends on glutamate and KtrA-KtraB potassium transporter. a) Glutamate is important for the extent of *Ziapin2* modulation of membrane potential dynamics. The peak hyperpolarisation response to light in the media with and without glutamate. Data from two independent experiments. Each dot is average of >100 cells. b-c) Membrane potential change following light stimulation (blue) with b) *yugO* and c) *ktrAB* deletion strains. *yugO* does not impact the hyperpolarization observed upon light stimulation. Mean ± sem from three independent experiments. KtrA-KtraB potassium channel is involved in Ziapin2-induced membrane potential modulation, as its deletion eliminates the hyperpolarization observed upon exposure to 470 nm light.

Towards better understanding the biological machineries of the process, we utilised potassium channel deletion mutant strains. We first tested the *yugO* deletion strain because the potassium channel encoded by this gene is known to mediate biofilm electrical signalling^[13]^. YugO channel is structurally similar to the classic KcsA potassium channel with a TVGYG selectivity filter motif. The photo-stimulation microscopy experiment was conducted in the same way as the wild type. We first confirmed that the TMRM signal is stable over the course of our experiment without *Ziapin2*. With *Ziapin2*, the TMRM signal underwent a transient signal increase upon light stimulation, similar to the wild type (Figure 5b, see also Figure S6 for negative control). Surprisingly, these results suggest that YugO channel is dispensable for the light-triggered hyperpolarisation, in spite of its role in biofilm electrical signalling.

We next tested the mutant strain that lacks the genes encoding the high-affinity potassium channel KtrAB, which belongs to TrK/Ktr/HKT super family ^[33]^. The TMRM signal was less stable with this strain than the wildtype and showed gradual signal decay in our negative control experiments (Figure 5c, left panel). Upon exposure to 470 nm light, no significant change in membrane potential was observed (Figure 5c, right panel, and Figure S5a). These results strongly suggest that KtrAB potassium channel may play a role in the response dynamics.

Our understanding of *B. subtilis* ion channels is currently incomplete, and it is possible that *Ziapin2* isomerisation triggers opening of uncharacterised ion channels. To explore this possibility, we employed three ion channels blockers: namely, the potassium channel blocker tetraethylammonium (TEA), the calcium channel blocker Nirendipine, and the chloride channel blocker Indanyloxyacetic acid-94 (IAA-94). The wildtype cells were treated with an ion channel blocker for 1 hr before being used for photo-stimulation microscopy experiments. The results showed that, in the presence of *Ziapin2*, cells treated with TEA or nitrendipine showed a TMRM signal increase upon light exposure, as it would happen in the absence of blockers (Figure 6a and 6b). On the other hand, cells treated with IAA-94 did not show a transient signal rise upon light stimulation (Figures 6c and S5b). Instead, we observed a slow gradual hyperpolarisation which is likely unrelated to *Ziapin2* isomerisation as the condition without *Ziapin2* showed a similar pattern. Altogether, our results suggest that *Ziapin2* isomerisation causes gating of ion channels (Figure 6d). In other words, separate to biofilm electrical signalling which is mediate by YugO, bacterial membrane is equipped with a machinery that can produce a bioelectric response to a fast voltage changes by *Ziapin2* isomerisation.

**Figure 6.**
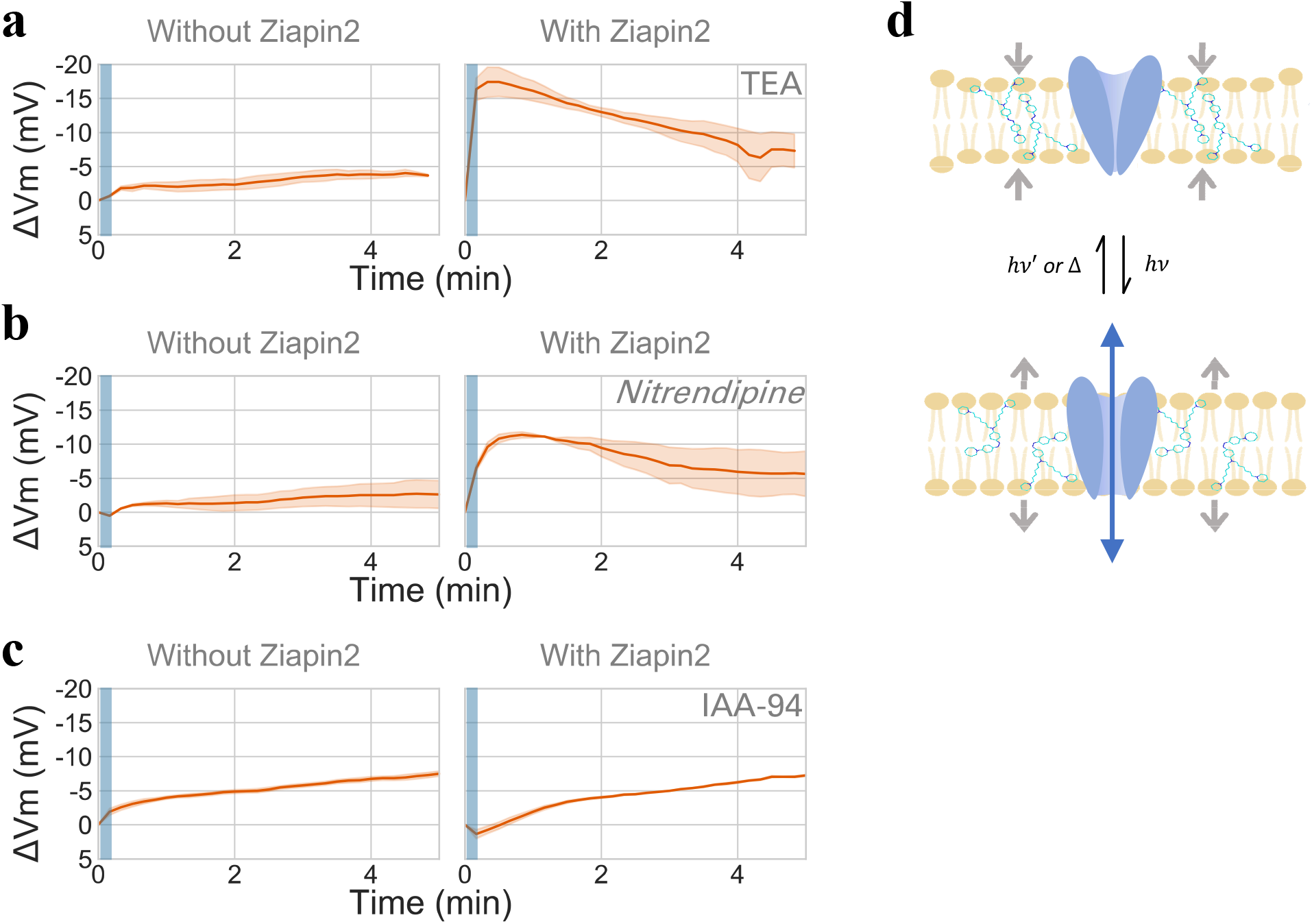
Chloride channel blocker attenuate the hyperpolarisation response. Membrane potential change over time in the presence of ion channel blockers, a) the potassium blocker TEA, b) the calcium blocker Nitrendipine, and c) the chloride blocker IAA-94. IAA-94 impairs the hyperpolarization induced by *Ziapin2* upon light stimulation, suggesting chloride channels are involved in *Ziapin2*-induced membrane potential dynamics. Mean ± sem from two independent experiments. d) Illustrative diagram showing opening of ion channels upon photo-induced *Ziapin2* isomerisation.

## Discussion

We demonstrate that the membrane potential of *B. subtilis* can be controlled by optostimulation without genetic modifications. To the best of our knowledge, this is the first example of inducing a transient membrane-potential dynamics using visible light. We employed a membrane-targeted azobenzene molecule, *Ziapin2*, which is able to drive modulation of the membrane capacitance and potential via an optomechanical effect. Under visible light illumination (λ ~ 470 nm), we observe a transient hyperpolarization followed by a depolarization rebound. The time-scale discrepancy between the relatively fast isomerisation process and the long-lasting biological effects prompted us to study the possible involvement of voltage-gated ion channels. Intriguingly, we found that the potential modulation brought about by *Ziapin2* isomerisation triggers the opening of the chloride channel, whose role is still largely uncharacterised for prokaryotes. More in general, this indicates that bacteria are equipped with bioelectric machinery that can respond to fast voltage changes. It is anticipated that future studies will further characterise the physiological roles of bacterial ion channels.

An important future research topic is elucidating the molecular mechanism of the bioelectric circuit. While cells exposed to the potassium channel blocker TEA exhibited photo-stimulated membrane potential dynamics, *ktrAB* deletion strain did not show such a response. The blockage by TEA depends on an aromatic residue on the extracellular side of the channel^[34]^, hence, it is possible that TEA does not block KtrAB channel. In a future project, we would also like to characterise the molecular identity of ion channels that are blocked by IAA-94. While many bacteria carry genes encoding chloride channels, which are commonly used as the model for neural ion channels, the physiological roles of chloride channels are still largely elusive. Our finding could be a ground to elucidate the physiological roles of chloride channels. Another important group of channels to investigate further is mechanosensitive channels^[35]^.

To date, the bioelectronics community’s efforts to interrogate cells have primarily been devoted to eukaryotes^[36–38]^, yet the community has recently steered to the development of new interfaces for studying and controlling bacterial functions^[12,24,39–41]^. The interest is mostly driven by the recent observation of neuron-like electrical patterns, such as spiking^[14]^ and oscillation^[13,42]^. It is intriguing to analogously consider these signalling and circuits as forming a “bacterial brain” that regulates metabolism and adaptation/responsivity to external stimulus and stressors, such as drugs and antibiotics. The fact that the bacterial membrane potential can be dynamically controlled by external stimuli opens new and exciting opportunities to gain new biological insights connected to signalling roles of the bacterial membrane potential. Exogenous light stimulation is perfectly suited to serve to this role, as it permits to elicit signalling with high spatiotemporal precision and remotely, therefore surpassing some intrinsic limitation of electrode-based methods, such as the need for contacting small, motile and highly heterogeneous bacterial cells.^[43]^

For these reasons, non-genetic optostimulation has the potential to boost research in the field of bacterial electrophysiology, for instance via the use of patterned optical excitation/probing at different nodes of the neuron-like network, as well as to facilitate the development of new synthetic-biology technologies for the bioelectrical engineering of bacterial functions.

## Material and Methods

### Synthesis of Ziapin2

*Ziapin2* has been synthesised according to the procedure that has been already published.^[10,11]^ Unless otherwise stated, all chemicals and solvent were commercially available and used without further purification. Reactions of air- and water-sensitive reagents and intermediates were carried out in dried glassware and under argon atmosphere. If necessary, solvents were dried by means of conventional method and stored under argon. Thin layer chromatography (TLC) was performed by using silica gel on aluminium foil, Sigma Aldrich). NMR spectra were collected with a Bruker ARX400. Mass spectroscopy was carried out with a Bruker Esquire 3000 plus.

### Growth conditions and preparation of agarose pads

Glycerol stock of Bacillus subtilis NCIB 3610 wild-type strain (WT) was streaked on lysogeny-broth (LB) 1.5% agar and incubated overnight in a 37°C non-shaking incubator. A single colony was picked from this plate, inoculated in LB and incubated at 37°C shaking overnight. When specified in the text, a genetically modified strain (listed in Table S1) was used instead of WT. When culturing a strain with antibiotic-resistance genes, appropriate antibiotics were added to the media in the following concentrations: spectinomycin 100 μg/mL; kanamycin 5μg/mL. Following overnight cultivation in liquid LB, cells were pelleted and washed once with resuspension media (RM) ^[44]^ (RM; composition per1 litre: 46 μg FeCl2, 4.8 g MgSO4, 12.6 mg MnCl2, 535 mg NH4Cl, 106 mg Na2SO4, 68 mg KH2PO4, 96.5 mg NH4NO3, 219 mg CaCl2, 2 g monosodium L-glutamate), and then incubated in RM at 37°C shaking for an hour prior to microscopy assay. When specified in the text, glutamate was omitted from RM. Following incubation with RM, cells were then deposited on RM 1.5% weight/volume Low Melting Point (LMP) agarose pads prepared as described previously ^[19,22,23]^. When specified, TMRM, *Ziapin2* and ion channel blockers were added at the following concentrations: TMRM at 100 nM (Molecular Probes); *Ziapin2* at 1 μg/mL; TEA (Sigma-Aldrich) at 25 mM; Nitrendipine (Sigma-Aldrich) at 10 μM; IAA-94 (ApexBio Technology) at 100 μM.

### Time-lapse microscopy and light stimulation

For time-lapse and 470 nm light stimulation experiments, the fluorescence microscope Leica DMi8, equipped with an automated stage, Hamamatsu Orca-flash 4.0 scientific CMOS (complementary metal–oxide–semiconductor) camera, a PeCon incubation system, and an objective lens HCX PL FLUOTAR 100x/1.30 OIL PH3, was used. TMRM fluorescence was detected with 500 ms exposure with Ex554/23 and Em609/54 filters (Semrock). The white LED of SOLA-SM II light engine (Lumencor) was used with the power level 10/255 (~4% of full power). For 470 nm stimulation Ex466/40 filter (Semrock) was used with 10 seconds exposure, and when specified in the text, the power level of the white LED of SOLA-SM II light engine was varied from 2/255 to 10/255. The light power of the 470 nm stimulation was measured with the PM16-121 power meter (Thorlabs) and the power density calculated in accordance with the area of the field of view.

Time-lapse duration was 2 minutes before 470 nm stimulation, with acquisition interval of 10 seconds. Immediately after, another 5 minutes time-lapse with same acquisition interval was conducted, where 470 nm exposure occurred once after the first TMRM image acquisition.

### Membrane potential estimation

Estimation of *B. subtilis* membrane potential changes (Δ*v_m_*) from the fluoresce intensity was performed as described by Ehrenberg *et al*.^[32]^ using the following equation:

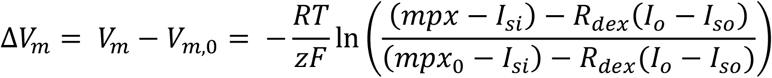

where *V_m_* is membrane potential, *V*_*m*,0_ is the resting membrane potential, *R* is the gas constant, *T* is the temperature in Kelvin, *z* is the charge of the dye, *F* is the Faraday constant, *mpx* is the mean pixel intensity from analysed cells, *mpx*_0_ is the mean pixel intensity of cells before light stimulation, *I_o_* is the mean background intensity, *I_si_* is the autofluorescence of the cell (measured from cells without TMRM) and *I_so_* is the background autofluorescence in the absence of TMRM. *R_dex_* accounts for off-focus signal. For our experimental setup, *R_dex_* was determined to be 0.976 by taking the ratio of off-focus and in-focus image with rhodamine dextran as described by Ehrenberg *et al*.^[32]^. Calculations were performed with JupyterLab 1.2.6 ^[45]^.

### Steady-stated UV-Vis/PL spectroscopy and ζ potential measurements

Cells were suspended in PBS to OD_600nm_ = 0.5. For ζ potential measurements, 100 mL of each sample was diluted into 900 mL PBS. The measurements were performed on a Malvern Zetasizer Nano ZS (Malvern Instruments, Malvern, U.K.) at RT. Data points given are an average of 3 biological replicates with 3 measurements each.

UV-Vis absorption measurements were performed using a Perkin Elmer Lambda 1050 spectrophotometer, with deuterium (180–320 nm) and tungsten (320–3300 nm) lamps, a monochromator and three detectors (photomultiplier 180–860 nm, InGaAs 860–1300 nm, and PbS 1300–3300 nm). Absorption spectra were normalized according to a reference spectrum taken at 100% transmission (without the sample), 0% transmission (with an internal shutter), and in the presence of the reference solvent. For the PL measurements and the excitation profiles an iHR320Horiba NanoLog Fluorometer was employed, equipped with a Xenon lamp, two monochromators, and two detectors (photomultiplier and InGaAs).

### Ziapin2 cellular uptake experiments

Cells suspended in PBS were stained with different concentrations of *Ziapin2* and kept at 37°C for 60 minutes in dark. The samples were then centrifuged and 200 μl of each supernatant was transferred to a clean 96-well plate for UV-Vis absorption with a Tecan Spark10M plate reader. The light excited samples (LED 470 nm) were treated using the following illumination protocol: 10 minutes of light followed by 10 minutes in dark conditions, repeated three times. Absorbance was measured at 490 nm. Control samples with no cells were treated the same, and their absorbance values represented the total molecule for reference. All conditions and controls were measured in triplicate.

### Time-resolved PL measurements

TRPL experiments were carried out using a femtosecond laser source coupled to a streak camera detection system (Hamamatsu C5680). A Ti:sapphire laser (Coherent Chameleon Ultra II, pulse bandwidths of B140 fs, repetition rate of 80 MHz, and maximum pulse energy of 50 nJ) was used to pump a second-harmonic crystal (b-barium borate) to tune the pump wavelength to 470 nm. The measurements here shown were performed recording the first 130 ps of decays, with an IRF of 4.1 ps. When required, a Peltier cell was used in order to control the temperature of the sample.

## Supporting information

Supporting Information is available from the Wiley Online Library or from the author.

## Acknowledgments

We thank Dr Fabio dos Santos Rodrigues and Pietro Bertolotti for their comments on the manuscript. G.M.P. acknowledges the financial support from Fondazione Cariplo, grant no. 2018-0979. TDS and MA acknowledge the Biotechnology and Biological Sciences Research Council (BBSRC)/Engineering and Physical Sciences Research Council (EPSRC) grant to the Warwick Integrative Synthetic Biology Centre (Grant BB/M017982/1).

## Data Availability Statement

The data that support the findings of this study are available from the corresponding authors upon reasonable request.

