## Supplementary material for "Membrane Targeted Azobenzene Drives Optical Modulation of Bacterial Membrane Potential": SI

[1]School of Life Sciences, University of Warwick, Coventry, CV4 7AL, UK
[2]Center for Nanoscience and Technology, Istituto Italiano di Teconologia, Milano, 20133, Italy
[3]Physical and Theoretical Chemistry Laboratory, Oxford, OX1 3QZ, United Kingdom
[5]Department of Chemistry. Materials and Chemical Engineering "Giulio Natta" Politecnico di Milano, Milano, 20133, Italy
[5]Department of Physics, Politecnico di Milano, Milano, 20133, Italy

#These authors contributed equally to this work

### Supplementary data

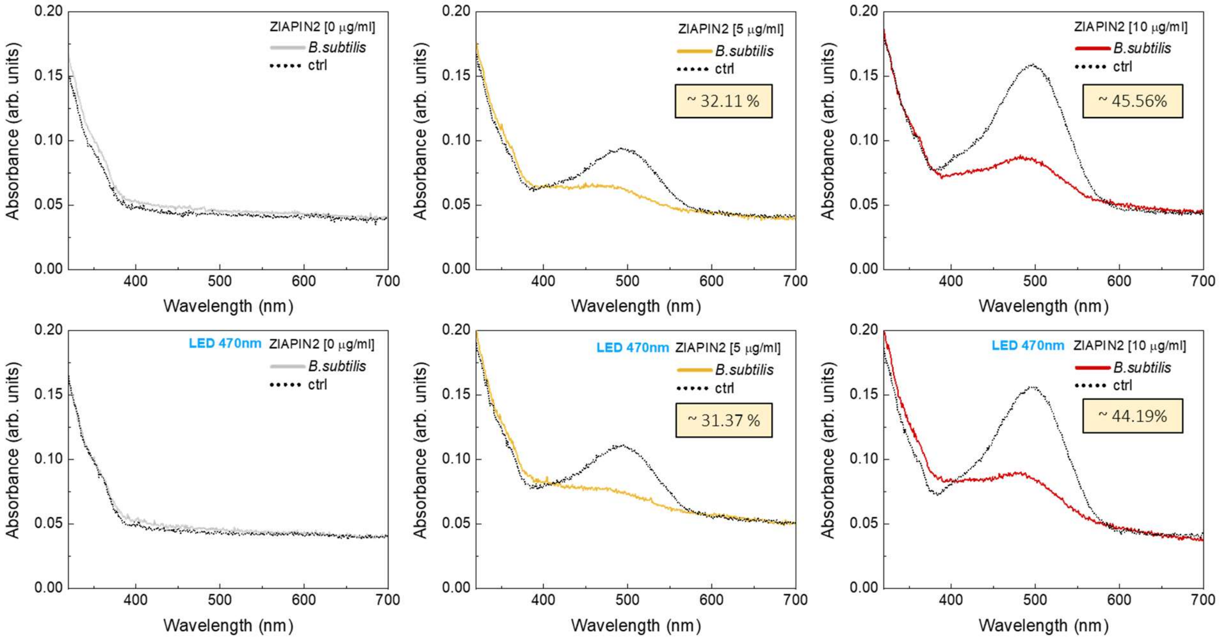


**Figure S1.** Cellular uptake experiments performed for 0, 5 and 10 µg/mL of Ziapin2, in the supernatant (dashed line) and in the cell fraction (continuous line). We carried out the experiment in dark and under illumination (470 nm), showing that light excitation does not cause ZIAPIN2 detachment from cells

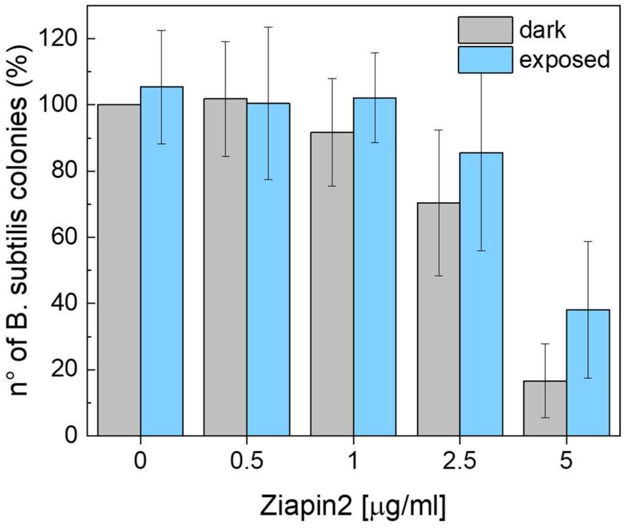


**Figure S2.** *B. subtilis* cells viability upon administration of Ziapin2 assessed via plate reader assay under dark and light (470 nm) conditions.

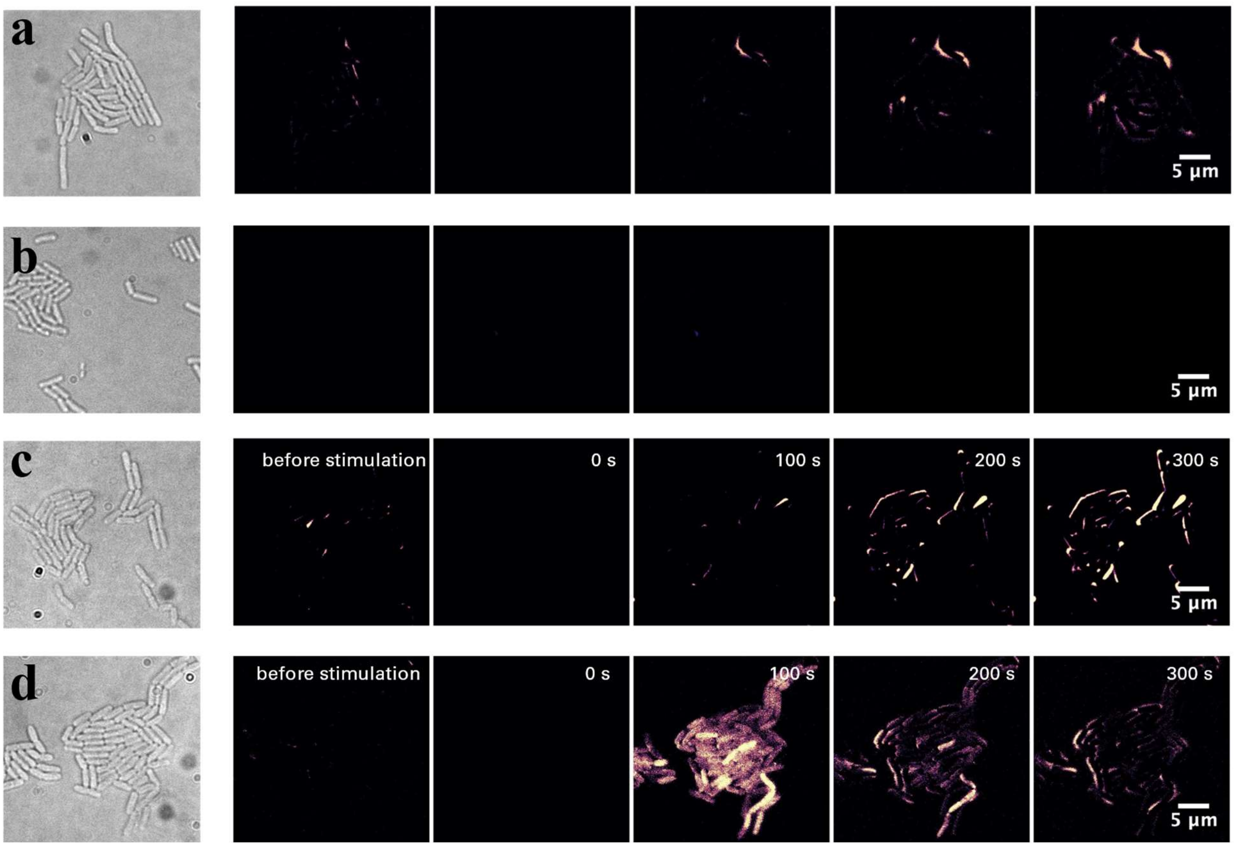


**Figure S3.** Images of *B. subtilis* membrane potential overtime. **a**) Cell exposed to TMRM only, not stimulated with light – no hyperpolarization observed. **b**) Cell exposed to TMRM and Ziapin 2, not stimulated with light – no hyperpolarization observed. **c**) Cell exposed to TMRM only, stimulated with 470nm light for 10 seconds – no hyperpolarization observed. **d**) Cell exposed to TMRM and Ziapin2, stimulated with 470nm light for 10 seconds – hyperpolarization is observed, followed by depolarization.

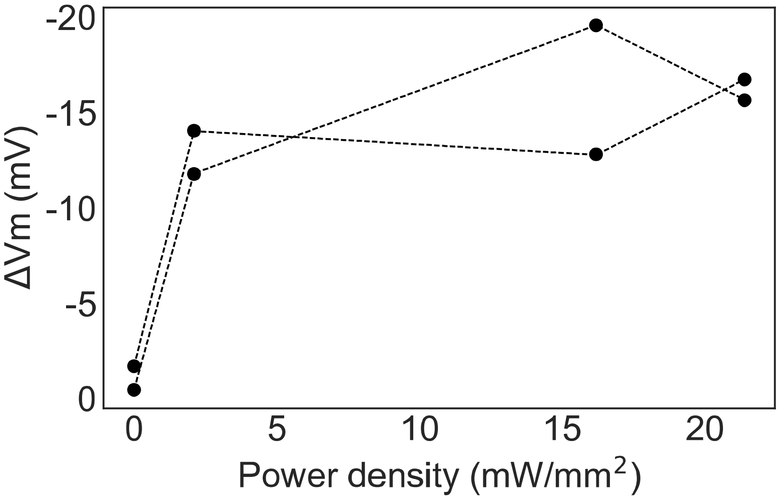


**Figure S4.** 470 nm light power density of 2 mW/mm$^2$ is enough to induce Ziapin2-dependent modulation of membrane potential. A peak of induced hyperpolarization is observed with 16 mW/mm$^2$.

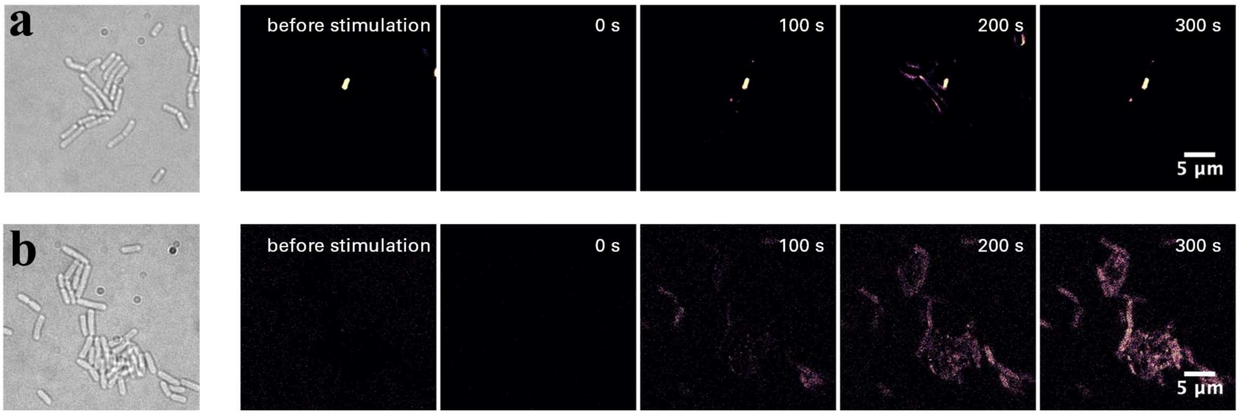


**Figure S5.** The potassium channel ktrAB and chloride ions are involved in the Ziapin2 induced membrane potential modulation. **a**) ΔktrAB cells exposed to TMRM and Ziapin2 do not show hyperpolarization upon 470 nm light stimulation. **b**) Wildtype cells exposed to TMRM, Ziapin2 and 100 µM of chloride blocker IAA-94 also do not show hyperpolarization upon 470 nm light stimulation.

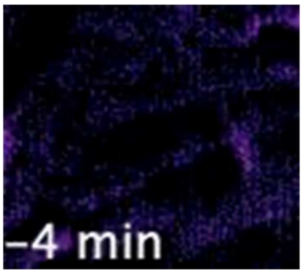


**Movie S1.** *B. subtilis c*ells exposed to TMRM and Ziapin2. After 470nm light stimulation, cells are hyperpolarized, and gradually depolarize overtime.

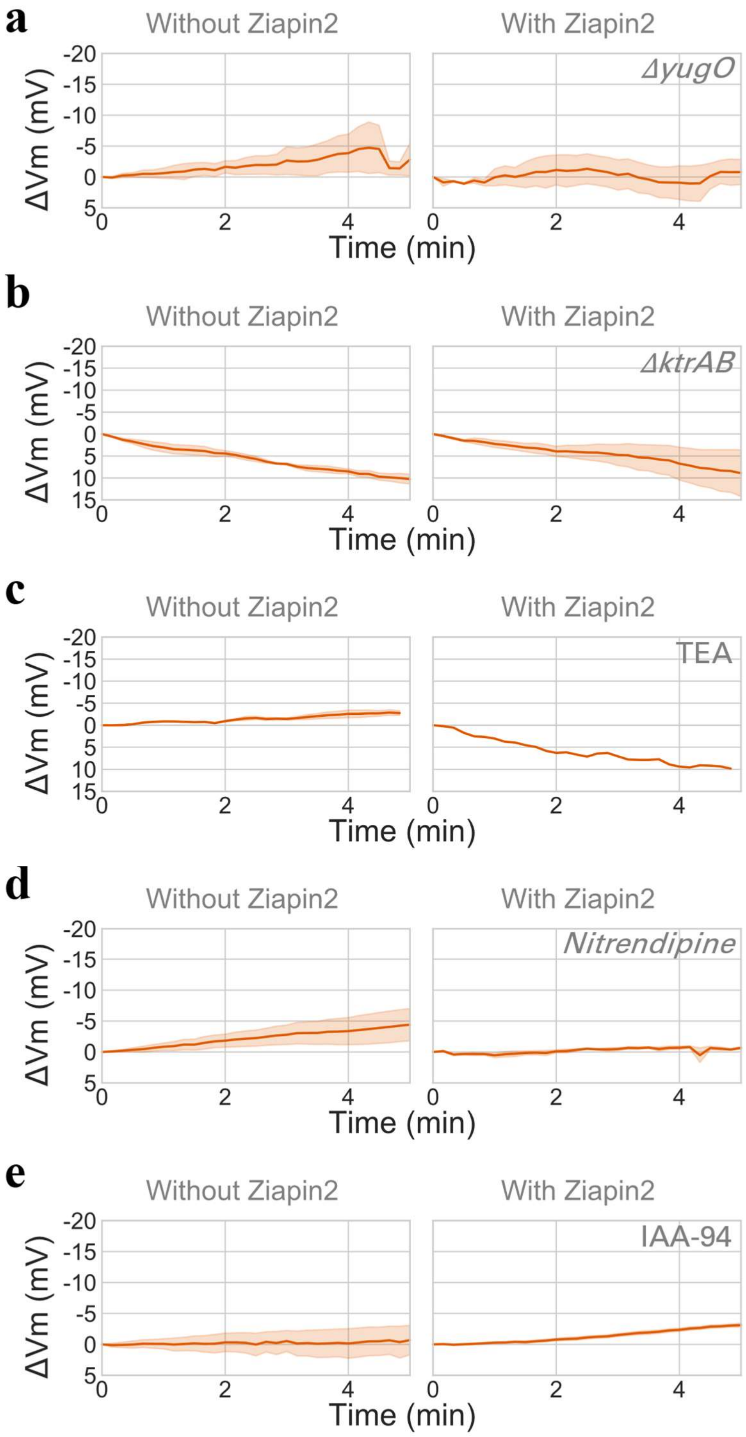


**Figure S6 –** No 466nm light-stimulation controls. **a**) Δ*yugO* cells with and without Ziapin2. **b**) Δ*ktrAB* cells with and without Ziapin2. **c**) WT cells exposed to TEA, with and without Ziapin2. **d**) WT cells exposed to Nitrendipine, with and without Ziapin2. **d**) WT cells exposed to IAA-94, with and without Ziapin2.

**Table S1**

| *B. subtilis* strains | | |
|---|---|---|
| **Strain** | **Genotype** | **Source** |
| NCIB3610 | Wildtype | Gift from Süel lab |
| PY79 | Wildtype | Gift from Süel lab |
| Δ*ktrAB* | ktrAB::neo | Bacillus Genetic Stock Centre (https://bgsc.org/) |
